# Systems Biology under heat stress in Indian Cattle

**DOI:** 10.1101/2020.04.09.031153

**Authors:** Raja Ishaq Nabi Khan, Amit Ranjan Sahu, Waseem Akram Malla, Manas Ranjan Praharaj, Neelima Hosamani, Shakti Kumar, Smita Gupta, Shweta Sharma, Archana Saxena, Anshul Varshney, Pragya Singh, Vinay Verma, Puneet Kumar, Gyanendra Singh, Aruna Pandey, Shikha Saxena, Ravi Kumar Gandham, Ashok Kumar Tiwari

## Abstract

Transcriptome profiling of Vrindavani and Tharparkar cattle revealed that more numbers of genes dysregulated in Vrindavani than in Tharparkar. A contrast in gene expression was observed with 18.5 % of upregulated genes in Vrindavani were downregulated in Tharparkar and 17.5% upregulated genes in Tharparkar were downregulated in Vrindavani. Functional annotation of genes differentially expressed in Tharparkar and Vrindavani revealed that the systems biology in Tharparkar is moving towards counteracting the effects due to heat stress. Unlike Vrindavani, Tharparkar is not only endowed with higher expression of the scavengers (*UBE2G1, UBE2S*, and *UBE2H*) of misfolded proteins but also with protectors (*VCP, Serp1*, and *CALR*) of naïve unfolded proteins. Further, higher expression of the antioxidants in Tharparkar enables it to cope up with higher levels of free radicals generated as a result of heat stress. In this study we found relevant genes dysregulated in Tharparkar in the direction that can counter heat stress.

## Introduction

Cattle being homoeothermic modulate their internal body temperature in sync to environmental temperature by equilibrating the amount of heat produced within the body and dissipating it to the ambient environment. The stress that arises due to disproportionate thermodynamic behavior between cattle and its surrounding environment is termed as heat stress ^1^. Environmental induced hyperthermic stress lowers feed intake, which in turn reduces growth, milk production and reproductive efficiency, thereby negatively affecting the economics of livestock keepers ^2-4^. Heat stress has been associated with reduced fertility through its deleterious impact on oocyte maturation and early embryonic development ^5^. Increased morbidity and mortality was observed in animals due to immune depressive effect of heat stress ^6^.

India has a wide variety of indigenous cattle breeds distributed throughout its agro-climatic zones. These are known for their natural tolerance to tropical heat ^7,8^.To meet the growing demand for milk and to combine the heat tolerance and tick resistance of zebu with the productivity of temperate dairy breeds ^9^ several crossbreeding programs were taken up in India. Every state had its own crossbreeding policy, which is agro-climatic and breed-specific. Though the zebu crosses with European breeds produced more milk than zebu, they were found not withstanding heat/solar radiation ^10^. Crossbreds are susceptible to tropical diseases and require a constant input of good management conditions ^8^. Antagonism exists between heat tolerance and milk productivity ^11^. The adaptive capacity to heat stress varies between species and genetic groups within species. Among various adaptive mechanisms, physiological adaptability seems to be the primary step in cattle. Sahiwal cows better regulate body temperature in response to heat stress than Karan Fries ^8^. It was observed that Ongole cattle rely on the respiration rate to maintain thermal balance, while, Bali cattle rely on rectal temperature ^12^. In Brazil, Sindhi and Girolando breeds showed better physiological response to thermal stress than Gir cattle ^13^. Increase in respiration rate was reported in Nellore breed when exposed to heat load ^14^.

In India, Tharparkar is one among the best dairy breeds. It is adapted to the Indian states of Punjab and Haryana. ^15,16^. It is considered to be the hardiest, disease resistant, heat tolerant and tick resistant indigenous cattle breed of the country ^17^. This breed has also been used in several crossbreeding programs. Currently, percentage of purebreds is exceptionally low in India (Department of Animal Husbandry & Dairying, Govt. of India). Most of the farmer’s in India have Crossbreds and the percentage of exotic inheritance in these Crossbreds is unknown. Vrindavani, a Crossbred (synthetic), with 27% of Indigenous blood and 73% of exotic inheritance ^18^ is a representation of the kind of admixture that prevails in Indian cattle. Therefore, comparing Tharparkar with Vrindavani may establish the lost importance of Indigenous cattle and would further emphasize the need of conserving our indigenous purebreds because of advantageous traits like Heat tolerance.

Studies explaining the difference between the genetic groups (Crossbreds and Indigenous cattle) have been done mainly to address the physiological responses vis – a – vis heat stress and very few studies at the genomic level have been taken up ^19,20^.Also, there is an increasing need to develop methods by combining the knowledge from -omics technologies to identify heat tolerant animals ^11^. Transcriptome profiling / RNA-Sequencing (RNA-seq) is a high throughput omics approach to measure relative global changes in the transcripts under specific condition(s) ^21-23^ to study the systems biology behind a phenotype ^23,24^. RNA - seq allows for analysis of transcriptome in an unbiased way, with, a tremendous dynamic detection range (>8,000 fold), and low background signals ^25^. It has been used as an investigating tool in understanding disease pathogenesis ^26,27^ and differential physiological response to various biotic and abiotic factors ^28,29^.

In this study, Tharparkar and Vrindavani cattle were subjected to heat stress and blood samples were collected on 0^th^ day and 7^th^ day, as it is known that short term acclimation occurs around 5 − 6 days ^30,31^. The transcriptome of 7^th^ day was compared and with 0^th^ day in both the genetic groups to understand their differential response to heat stress.

## Results

### Physiological Parameters

The overview of the analysis is given in Figure 1. Respiration rate (RR), rectal temperature (RT) and T3 level increased significantly (p<0.05) on 7^th^ - day post heat stress in both the genetic groups (n=5) (Figure 2). However, the increase in RR, RT and T3 level, was found significantly (P<0.05) higher in Vrindavani than in Tharparkar.

**Figure 1:**
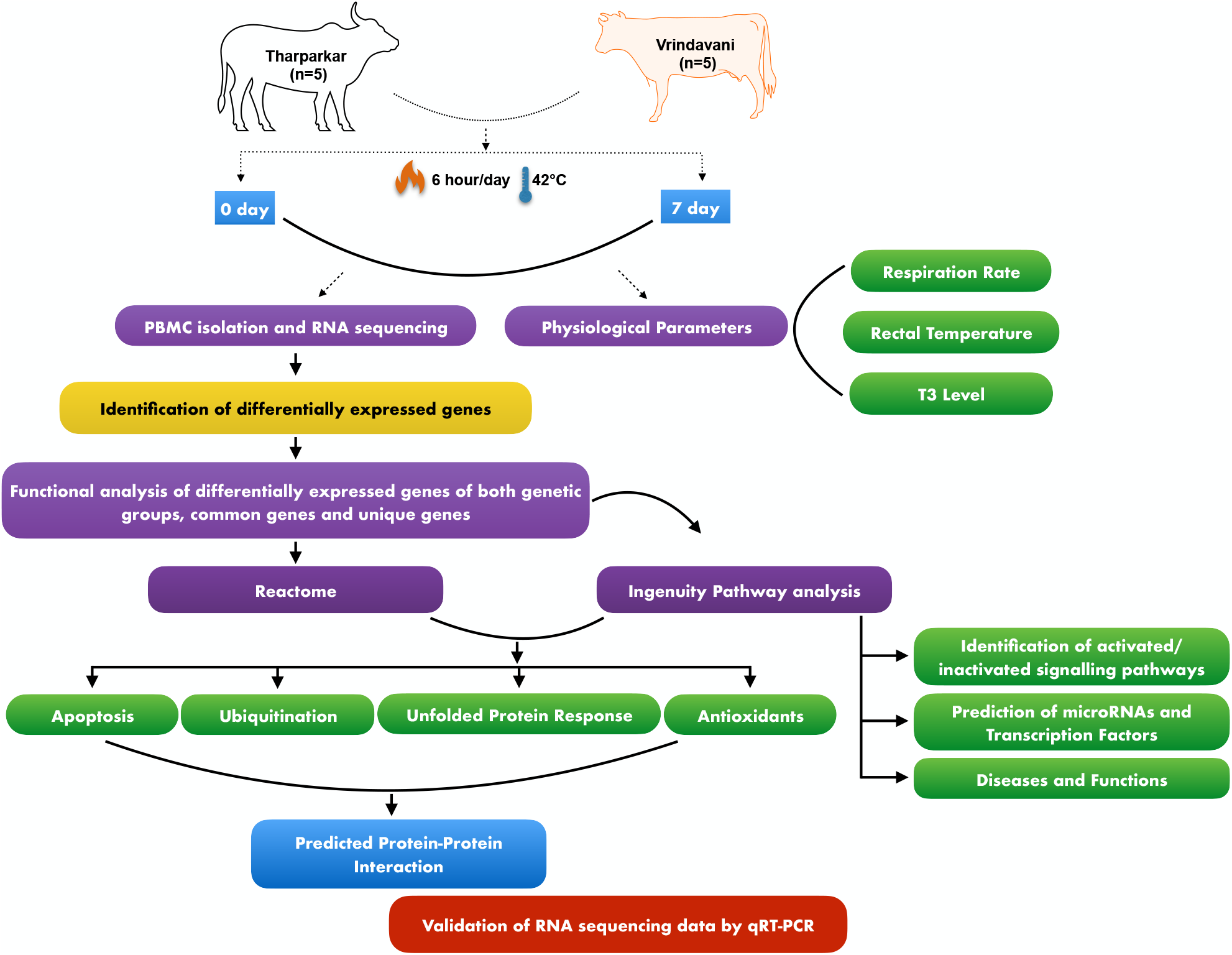
Overview of the work done: Two genetic groups (Tharparkar and Vrindavani) of cattle were exposed to a temperature of 42 °C for 7 days. Heat stress indicators - Respiration rate (RR), Rectal temperature and T3 level before exposure to heat (0day – control group) and at 7^th^ day of exposure (treated) were measured to evaluate heat stress. At these time points, RNA was isolated from PBMCs for high throughput sequencing. Transcriptome analysis was done to identify differentially expressed genes (DEGs) under heat treatment in both genetic groups. Genes involved in physiological processes (heat stress response, apoptosis, ubiquitination, unfolded protein response and antioxidant level) that are commonly associated with heat stress were compared between the two genetic groups. Further, functional annotation of DEGs was done using IPA.

**Figure 2:**
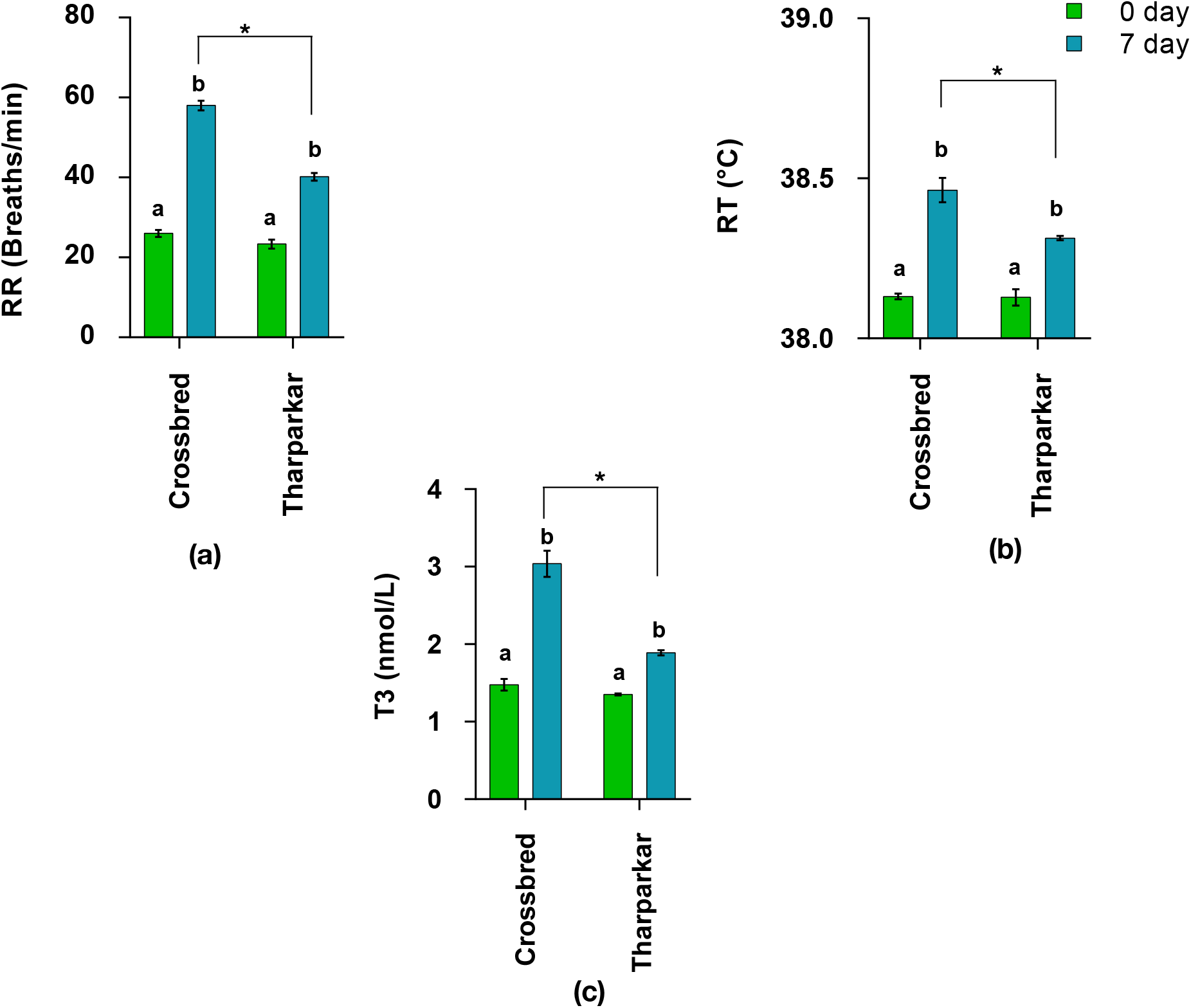
Respiration rate, Rectal Temperature and T3 level measured at 0 day (control) and 7 day post-heat exposure (treated) in Vrindavani and Tharparkar (n=5) Levels sharing the same superscript are not significantly (P > 0.05) different from each other.

### Comparison of DEGs of Vrindavani and Tharparkar under heat stress

The differentially expressed genes for each genetic group were obtained on comparing the 0 day and 7^th^ day RNA-seq data using EdgeR after obtaining the gene counts from RSEM. Under heat stress, global expression profiles of Vrindavani and Tharparkar were identified with 6042 and 4718 differentially expressed genes (DEGs), respectively (Supplementary Table 1). Among these, 3481 DEGs were found common between the two genetic groups, while 2561 and 1238 DEGs were uniquely found in Vrindavani and Tharparkar, respectively (Figure 3a). Additionally, 3132 and 2924 genes were upregulated and downregulated in Vrindavani, respectively, while 2367 and 2358 genes were upregulated and downregulated in Tharparkar, respectively (Figure 3b). On comparison of upregulated and downregulated genes, 724 and 1416 genes were found uniquely upregulated and 514 and 1145 genes were found uniquely downregulated in Tharparkar and Vrindavani, respectively. The comparison also revealed that 17.5% of upregulated genes (1278) in Tharparkar were downregulated in Vrindavani and 18.5% downregulated genes (1344) in Tharparkar were upregulated in Vrindavani. However, the number of common upregulated and downregulated genes in both the genetic groups were 357 (4.9%) and 498 (6.8%), respectively (Figure 3c).

**Figure 3:**
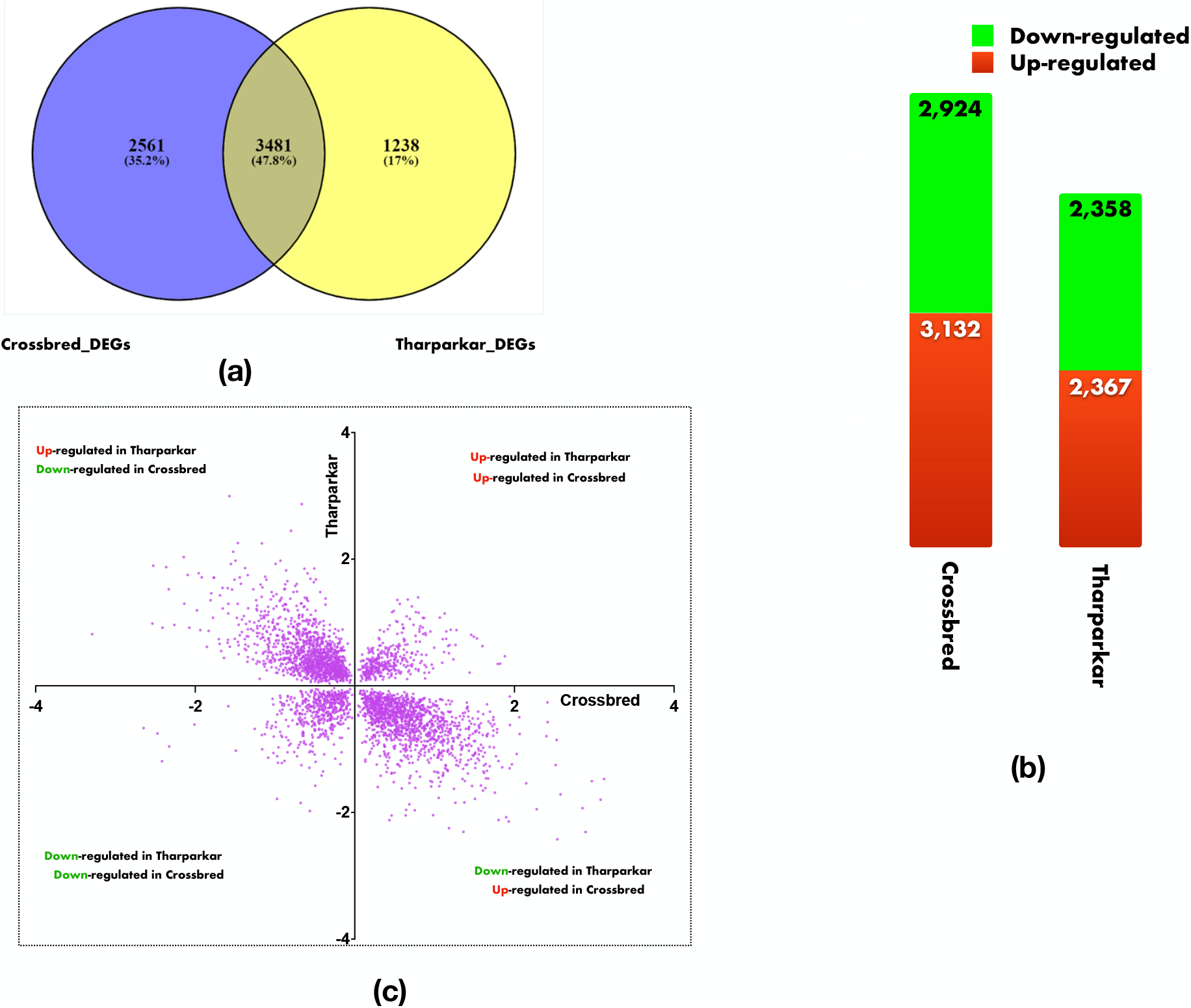
Expression of DEGs in Vrindavani and Tharparkar under heat stress: (a) Venn diagrams showing unique/common DEGs between Vrindavani and Tharparkar (b) Number of upregulated and downregulated in both genetic groups (c) Contrast in the expression of common DEGs

### Functional annotation of DEGs in Reactome

#### DEGs in Tharparkar and Vrindavani

On functional annotation of all the DEGs (6042 and 4718 in Vrindavani and Tharparkar, respectively), the commonly enriched significant pathways in both genetic groups included, neutrophil degranulation, L13a-mediated translational silencing of ceruloplasmin expression, Formation of a pool of free 40S subunits, ubiquitination and proteasome degradation, SRP-dependent cotranslational protein targeting to membrane and nonsense-mediated decay and selenocysteine synthesis. Among the most enriched pathways, peptide chain elongation, gap-filling DNA repair synthesis and ligation in TC-NER were the pathways significantly enriched in Tharparkar and not in Vrindavani. The pathways - SCF-beta-TrCP mediated degradation of Emi1, Autodegradation of Cdh1 by Cdh1:APC/C and Regulation of RAS by GAPs were the pathways significantly enriched in Vrindavani and not in Tharparkar.

#### Common DEGs in Tharparkar and Vrindavani

On functional annotation of common the DEGs (3481), the enriched significant pathways included, neutrophil degranulation, L13a-mediated translational silencing of ceruloplasmin expression, formation of a pool of free 40S subunits, ubiquitination and proteasome degradation, SRP-dependent cotranslational protein targeting to membrane, RNA Polymerase II Transcription Termination, Peptide chain elongation, nonsense-mediated decay and selenocysteine synthesis. On functionally annotation of 1278 common genes that were upregulated in Tharparkar and downregulated in Vrindavani, the enriched pathways included all the pathways as mentioned for the common except for ubiquitination & proteasome degradation. In addition, cellular response to stress was found significantly enriched these genes upregulated in Tharparkar and downregulated in Vrindavani. Further, on assessing the 1344 genes that were upregulated in Vrindavani and downregulated in Tharparkar, RUNX1 regulates transcription of genes involved in BCR signaling, p75NTR recruits signaling complexes, chromatin modifying enzymes, chromatin organization, death receptor signaling, resolution of D-loop structures through Holliday junction intermediates and TP53 regulates transcription of DNA repair genes pathways were significantly enriched.

#### Unique DEGs in Tharparkar

On functional annotation of unique DEGs (1238) expressed only in Tharparkar, the enriched significant pathways included, nucleotide excision repair, mitotic G1 phase and G1/S transition, DNA replication pre-initiation, cell cycle, synthesis of DNA, translation, DNA repair, activation of the pre-replicative complex, cell cycle checkpoints and inhibition of replication initiation of damaged DNA by RB1/E2F.

#### Unique DEGs in Vrindavani

On functional annotation of unique DEGs (2561) expressed only in Vrindavani, the enriched significant pathways included, unfolded protein response (UPR), loss of proteins required for interphase microtubule organization from the centrosome, amplification of signal from unattached kinetochores via a MAD2 inhibitory signal, mitotic spindle checkpoint, organelle biogenesis and maintenance and loss of Nlp from mitotic centrosomes.

### Functional annotation of DEGs in IPA

#### Canonical pathway analysis

Canonical pathway analysis by Ingenuity Pathway Analysis (IPA) revealed contrast in signaling pathways in Vrindavani and Tharparkar. Canonical pathways associated with Vrindavani and Tharparkar are represented in Figure 4a and 4b. In Vrindavani, Oncostatin M Signaling, Phospholipase C Signaling, EIF2 Signaling, Integrin Signaling, IL-3 Signaling, and CXCR4 Signaling were found to be highly inactivated (Z – score > 2.0) and PTEN signaling was found to be highly activated (Z – score < 2.0). In Tharparkar, EIF2 Signaling, Androgen Signaling, Oncostatin M Signaling, α-Adrenergic Signaling, BMP signaling pathway, and UVC-Induced MAPK Signaling were found to be highly activated and PTEN signaling was found to be inactivated. The canonical pathway Oncostatin M Signaling and EIF2 Signaling were found to have the highest ratio of genes involved vis-a-vis the genes in the database in Vrindavani and Tharparkar, respectively.

**Figure 4:**
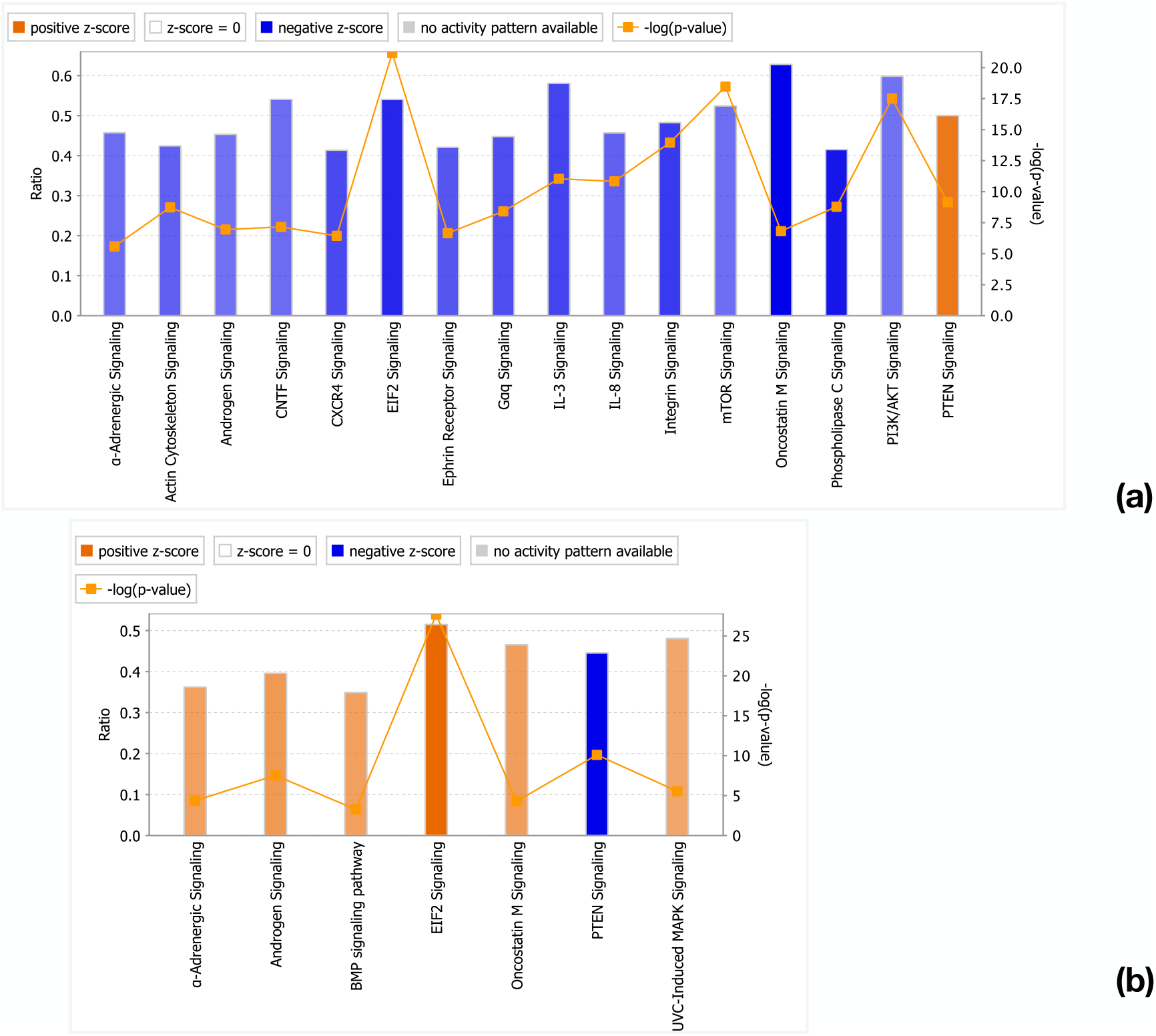
Canonical pathways activated/inactivated in (a) Vrindavani (b) Tharparkar under heat stress generated in the core analysis of Ingenuity pathway analysis tool. Orange color pathways are activated (Z > 2) and blue color pathways are inactivated (Z< −2). Height of the bar graphs indicates -log (p-value) and line graph showing the ratio of list genes found in each pathway over the total number of genes in that pathway.

While carrying out comparative analysis through IPA, Calcium-induced T Lymphocyte Apoptosis, BMP signaling pathway, UVC-Induced MAPK Signaling, Regulation of Cellular Mechanics by Calpain Protease, fMLP Signaling in Neutrophils, Melatonin Signaling, and Leukocyte Extravasation Signaling, were found inactivated in Vrindavani and activated in Tharparkar (Supplementary Figure 1). Genes involved in Oncostatin M Signaling-Growth factor receptor-bound protein 2 (*GRB2)*, GTPase HRas *(HRAS)*, Janus kinase 1 *(JAK1)*, Janus kinase 3 *(JAK3)*, Mitogen-activated protein kinase kinase 1 *(MAP2K1)*, Mitogen-activated protein kinase 1 *(MAPK1)*, Oncostatin-M *(OSM)*, Ras-related protein Rap-1b *(RAP1B)*, Ras-related protein Rap-2a *(RAP2A)*, Signal transducer and activator of transcription 1-alpha/beta *(STAT1)*, Signal transducer and activator of transcription 5B *(STAT5B)*, Non-receptor tyrosine-protein kinase *(TYK2)*, and Ras-related protein *(RRAS)* were found downregulated in Vrindavani and upregulated in Tharparkar (Figure 5a, b). While the key genes involved in PTEN Signaling pathway – Fas Ligand (*FASLG)*, member of RAS oncogene family *(RAP2A)*, Bcl-2-like protein 11 *(BIM)*,Caspase-3 *(CASP3)* and microspherule protein 1 (*MSP58)* were found upregulated in Vrindavani and downregulated in Tharparkar as well (Figure 6a, b).

**Figure 5:**
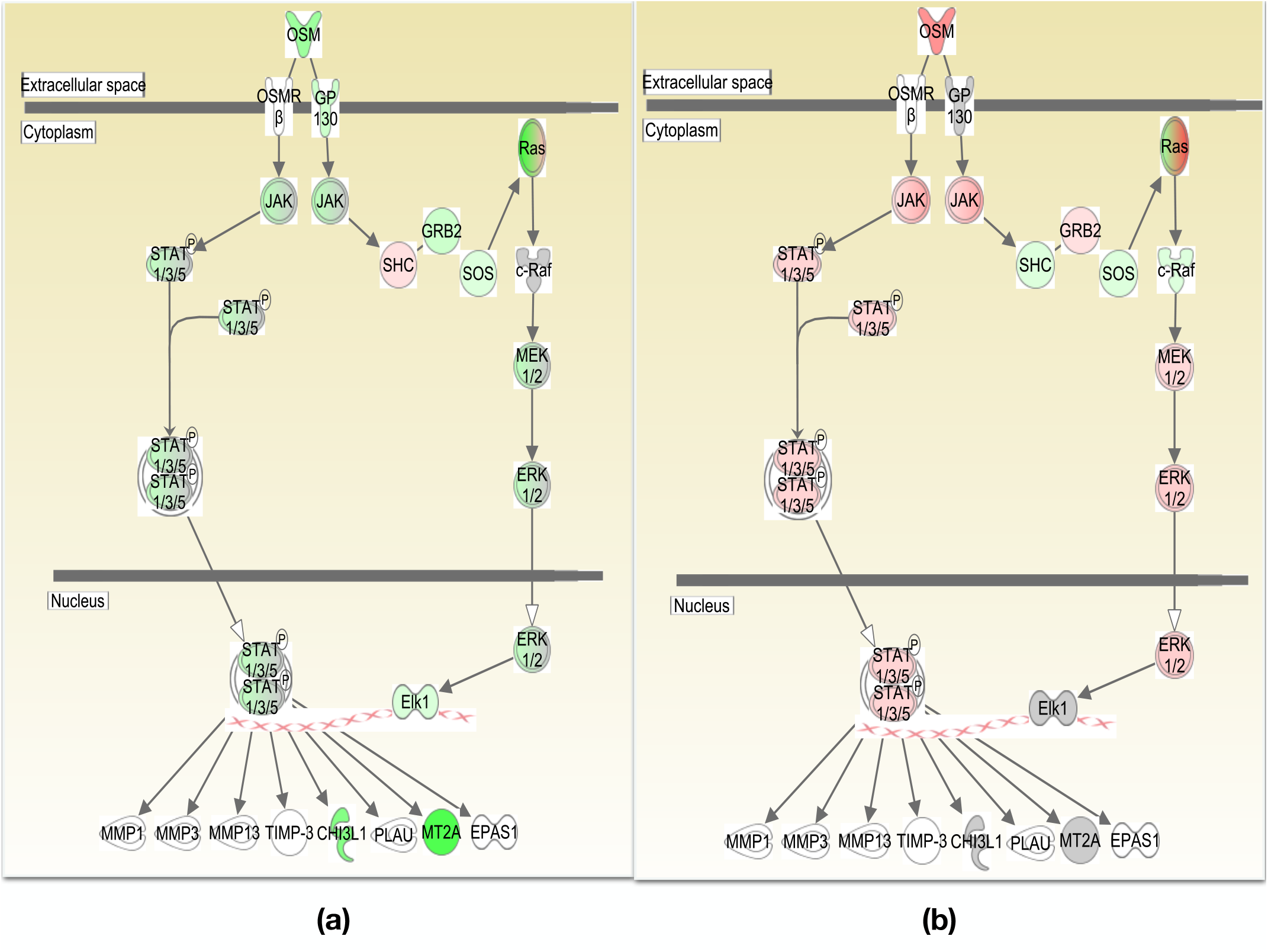
Canonical pathways generated in Ingenuity Pathway Analysis of Oncostatin M signaling pathway of DEGs in (A) Vrindavani, (B) Tharparkar. Genes that were upregulated are shown in red and downregulated in green. The intensity of red and green corresponds to an increase and decrease, respectively, in Log2 fold change. Genes in grey were not significantly dysregulated and those in white are not present in the dataset but have been incorporated in the network through the relationship with other molecules by IPA.

**Figure 6:**
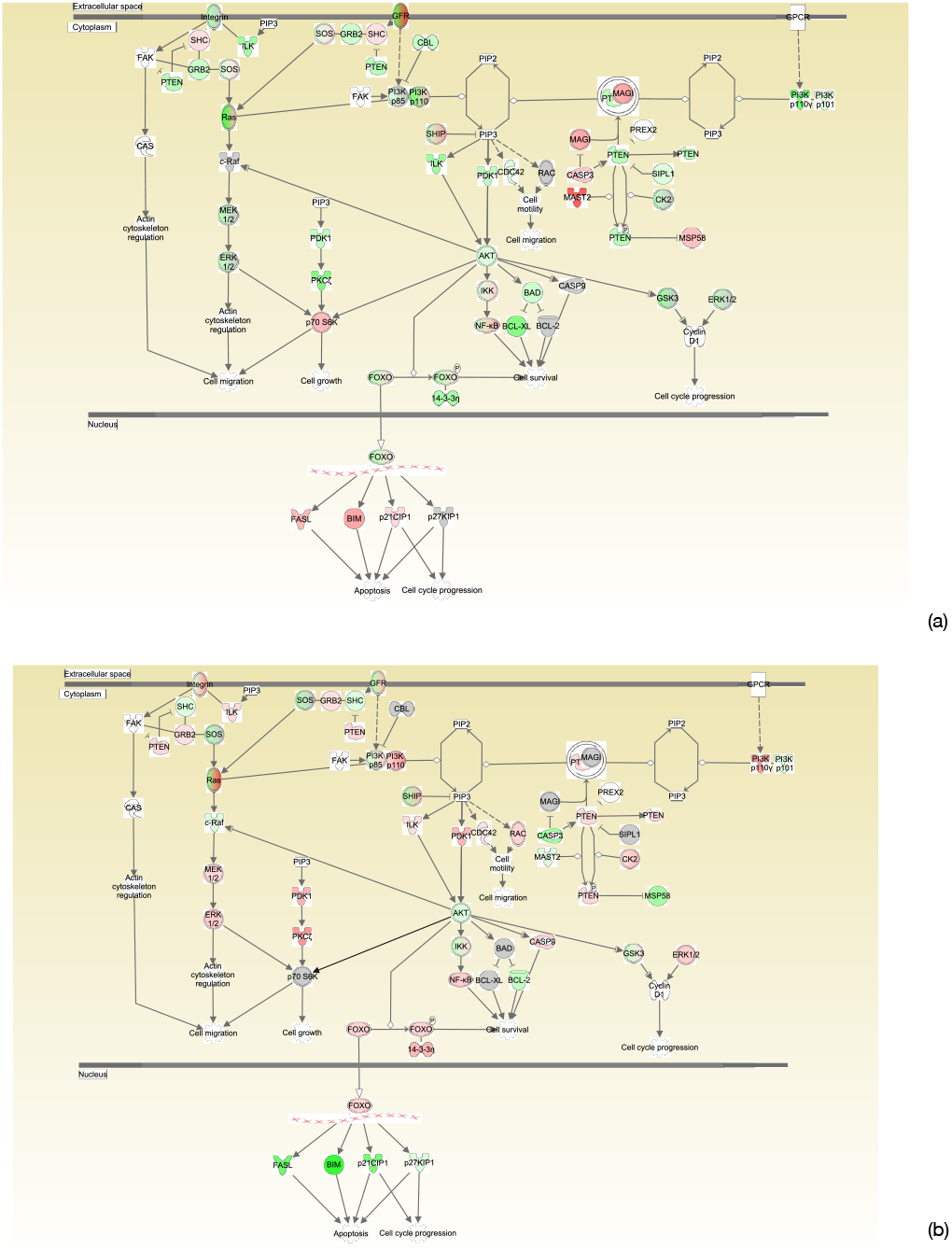
Canonical pathways generated in Ingenuity Pathway Analysis of PTEN signaling pathway of DEGs in (A) Vrindavani, (B) Tharparkar. Genes that were upregulated are shown in red and downregulated in green. The intensity of red and green corresponds to an increase and decrease, respectively, in Log2 fold change. Genes in grey were not significantly dysregulated and those in white are not present in the dataset but have been incorporated in the network through the relationship with other molecules by IPA.

#### Variation in microRNAs and Transcription factors

IPA, on evaluating the differentially expression genes predicts miRNAs and Transcription Factors (upstream regulators). In Vrindavani, 111 miRNAs were found to be inactivated and 37 activated. In Tharparkar, 205 miRNAs were found to be inactivated and 272 activated. Among them, 52 microRNAs were found common between the two genetic groups. Most of the common miRNAs were found activated in Vrindavani and inactivated in Tharparkar (Supplementary Figure 2). miR-4779, miR-4651, miR-1207-5p, miR-6967-5p and miR-504-3p are the top 5 miRNAs that were activated in Vrindavani and inactivated in Tharparkar.

Various Transcription factors were found to regulate the expression of the identified DEGs. Transcription factors, 19 in Tharparkar (11 activated and 8 inactivated) and 26 in Vrindavani (8 activated and 18 inactivated) were identified in IPA that regulate the expression of DEGs. Among them, *PAX5, MTA3, MYC, PROX1* and *SMAD7* in Vrindavani and, *HMGA1, MAF, MAX NOTCH22* and *NCOR1* in Tharparkar are the top 5 upregulated and activated TFs. On comparing the TFs of Tharparkar and Vrindavani, it was found that *BHLHE40, HMGA1, HMGB1, IKZF1*, and *TCF7* were found to be common. *BHLHE40, HMGA1*, and *TCF7* were found to be activated in Tharparkar and inactivated in Vrindavani and it was vice - versa with *HMGB1* and *IKZF1* (Supplementary Figure 3)

#### Disease and Functions

In the diseases and functions category, on evaluating all the DEGs in both the genetic groups, survival of the organism, ubiquitination, ubiquitination of protein and repair of DNA were found activated in Vrindavani and were found either inactivated/ not-activated in Tharparkar. Similarly, homeostasis, development of hematopoietic cells, leukopoiesis was found relatively activated in Tharparkar in comparison to Vrindavani. Further, translation, Expression of protein, translation of protein and degranulation were found inactivated in Vrindavani.

On correlating the results of disease and function with the results of reactome four major processes that were considered to be associated with heat stress viz. elicitation of unfolded protein response (UPR) in cells; Induction of apoptosis; Ubiquitination and; Imbalance in production of ROS and antioxidants. Heat shock genes and its associated genes are involved in elicitation of unfolded protein response (UPR) in cells. Heat shock genes have been found dysregulated under heat stress in both the genetic groups. Most of the genes encoding Heat shock proteins (HSPs) - Heat shock 70 kDa protein 4 (*HSPA4)*, Heat shock cognate 71 kDa protein *(HSPB8)*, Heat shock 70 kDa protein 1A *(HSPA1A)* ,Heat shock cognate 71 kDa protein *(HSPA8)*, Heat shock protein HSP 90-beta *(HSP90AB1)* and Heat shock protein HSP 90-alpha (*HSP90AA1*) and heat shock protein regulating factors-Heat shock factor 1 *(HSF1)* and Eukaryotic Translation Elongation Factor 1 Alpha 1 *(EEF1A1)* have been found to be downregulated/not-differentially expressed in Vrindavani but upregulated in Tharparkar. However, Calcium/Calmodulin Dependent Protein Kinase II Delta *(CAMK2D) that* is involved in the regulation of expression of heat shock genes was upregulated in Vrindavani and downregulated in Tharparkar.

Among the apoptotic genes, genes encoding Bcl-2-like protein 11 (*BCL2L11)*, Tumor necrosis factor ligand superfamily member 6 *(FASLG)*, TIR domain-containing adapter molecule 2 *(TICAM2)*, Toll-like receptor 4 *(TLR4)*, Adenomatous polyposis coli protein *(APC)*, Caspase-3 *(CASP3)*, Mitogen-activated protein kinase 8 *(MAPK8)*, Mixed lineage kinase domain-like protein *(MLKL)*, Late endosomal/lysosomal adaptor and MAPK and MTOR activator 5 *(XIP)*, Vimentin *(VIM)*, and High mobility group protein B2 *(HMGB2)* were found to be upregulated in Vrindavani and downregulated in Tharparkar. The number of upregulated genes involved in achieving the balance of ROS production and antioxidants, were found to be more in Tharparkar than in Vrindavani. Among these, genes encoding Glutathione peroxidase 3 (*GPX3)*, Nudix Hydrolase 2 *(NUDT2)*, Catalase *(CAT)*, Cytochrome c *(CYCS)*, Copper chaperone for superoxide dismutase *(CCS)*, Peroxiredoxin-5 *(PRDX5)*, Peroxiredoxin-6 *(PRDX6)*, Peroxiredoxin-1 *(PRDX1)*, Superoxide dismutase *(SOD1)*, and Cytochrome b-245 heavy chain *(CYBB*) were found either downregulated/not-differentially expressed in Vrindavani and upregulated in Tharparkar. More number of genes involved in Ubiquitination were differentially expressed in Vrindavani than in the Tharparkar. Genes encoding Ubiquitin-conjugating enzyme E2 G1 (*UBE2G1)*, Ubiquitin-conjugating enzyme E2 *(UBE2S)*, Ubiquitin-conjugating enzyme E2 H *(UBE2H)*, Ubiquitin A-52 residue ribosomal protein fusion product 1 *(UBA52)*, and Ubiquitin-activating enzyme E1 *(UBA1)* have been found downregulated/not-differentially expressed in Vrindavani and upregulated in Tharparkar. However, Valosin-containing protein *(VCP)*, RING finger protein 40 *(RNF40)*, and Ubiquitin-conjugating enzyme E2 L3 *(UBE2L3)* have been found downregulated in Vrindavani but not-differentially expressed in Tharparkar. Among the genes involved in Unfolded Protein folding response (UPR), genes encoding Membrane-bound transcription factor site-1 protease (*MBTPS1)*, Cyclic AMP-responsive element-binding protein 3-like protein 1 *(CREB3L1)*, Stress-associated endoplasmic reticulum protein 1 *(SERP1)*, Glycogen synthase kinase-3 alpha *(GSK3A)*, Eukaryotic translation initiation factor 2 subunit 3 *(EIF2S3)*, Calreticulin *(CALR)*, and Stress-associated endoplasmic reticulum protein 1 (*SERP1)* have been found downregulated in Vrindavani and upregulated in Tharparkar (Figure 7).

**Figure 7:**
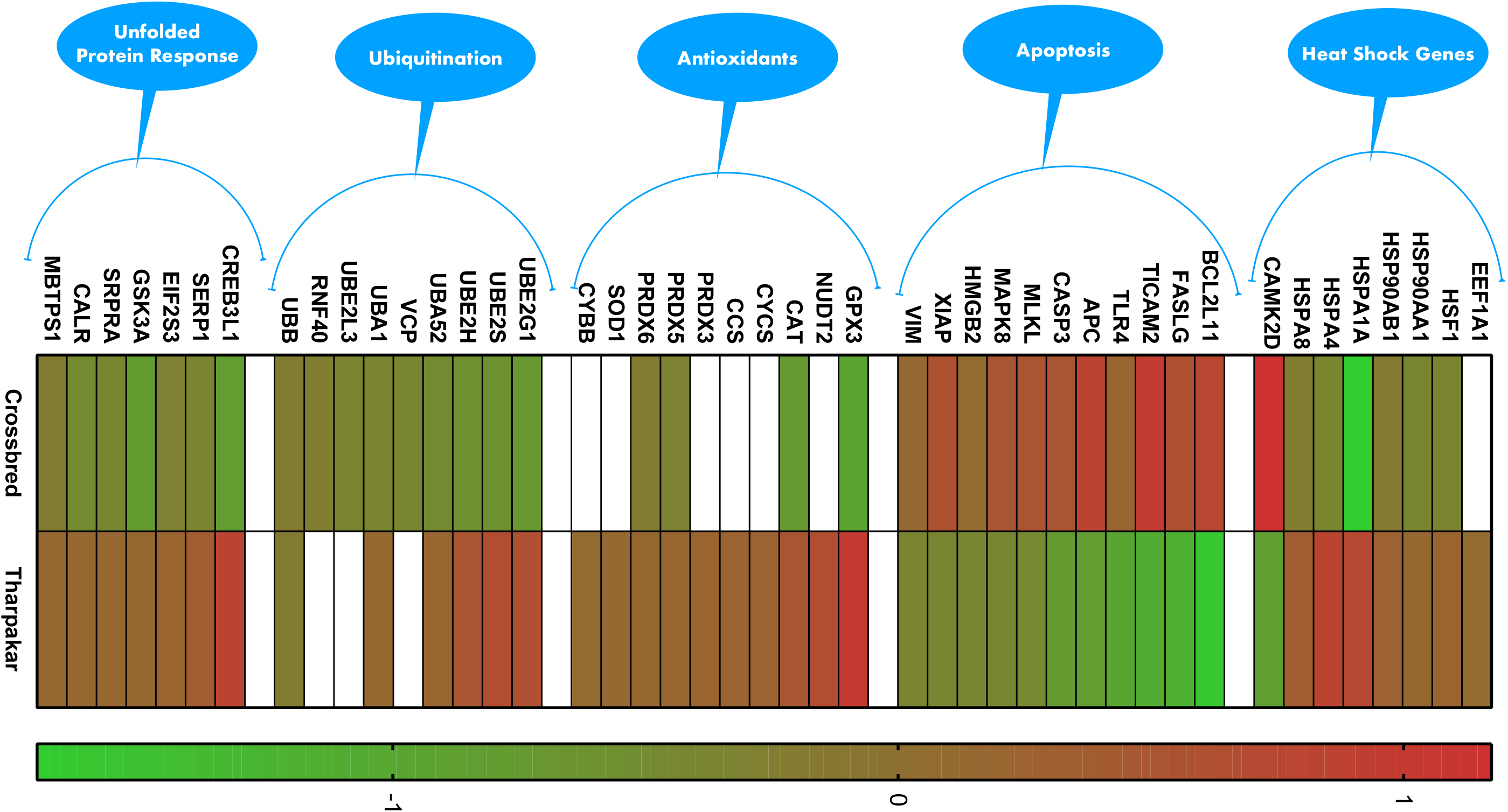
Contrast in the expression of genes involved in heat stress response, apoptosis, ubiquitination, unfolded protein response and balance in the production of ROS and antioxidants between two genetic groups.

### Protein - protein interaction (PPI) network

Among the genes involved in the 4 major processes considered, protein - protein interaction (PPI) network revealed functional importance of HSP70 (HSPA8 and HSPA1A) and ubiquitin (UBB, UBA52), in coordinating genes involved in heat stress. Out of the total 246 genes identified from the reactome database among these processes, 177 and 194 genes were found to be differentially expressed in Tharparkar and Vrindavani, respectively. Among these 126 genes were found to be commonly differentially expressed in Tharparkar and Vrindavani. PPI network for these common genes between Tharparkar and Vrindavani was constructed (Supplementary Figure 4). In PPI networks, hubs define the functional and structural importance of a network. The genes, which act as hubs in PPI networks were found to be *UBB, UBA52, HSPA8*, and *HSPA1A* (Supplementary Figure 4). Among the 4 hubs, *UBB* was downregulated in both genetic groups and the rest were downregulated in Vrindavani and upregulated in Tharparkar.

A change in the expression of the hub protein will have a larger effect than change in expression of non-hub proteins ^32^. Therefore, UBB, UBA52, HSPA8, and HSPA1A are taken to be critical for coordinating the changes in systems biology under heat stress. The hubs HSPA8 and HSPA1A are connected to genes that are associated with regulation of stress viz. nucleoporins genes - *NUP188, NUP155, NUP210* & *NUP214*; BAG family molecular chaperone regulators - *BAG1, BAG3* & *BAG4*; Heat Shock Protein Family A - *HSPA5, HSPA4, HSPA12B* & *HSPA9*; DnaJ Heat Shock Protein Family i.e. HSP40 - *DNAJA1, DNAJC2* & *DNAJB6*; Heat shock factor - *HSF1*; Ubiquitin - *UBB* & *UBA52* and; Sirtuin - *SIRT1*. The hubs - UBB and UBA52 are connected to molecules of different proteasome subunits viz. ⍰ type subunits - *PSMA1* & *PSMA2*; β type subunits - *PSMB4* & *PSMB8*; ATPase subunits - *PSMC2* & *PSMC5* and non-ATPase subunits - *PSMD2* & *PSMD13*. These hubs were also found connected to ubiquitin specific peptidases - *USP9X* and *USP7* and Ubiquitin-conjugating enzyme - *UBE2B, UBE2G1, UBE2Z, UBE2H, UBE2J2, UBE2S* & *UBE2D2*.

### Real-time validation

Six genes (*HSF1, SOD1, CALR, GSK3A, CAT* & *GPX3*) that were upregulated in Tharparkar but downregulated/not expressed in Vrindavani and four genes (*CASP3, FASLG, BCL2L11* & *APC*) that were upregulated in Vrindavani but downregulated in Tharparkar were considered for Real time PCR based on their role in heat stress. The expression of genes was in concordance with the RNA-Seq results (Supplementary Figure 5 and Supplementary table 2).

## Discussion

Heat stress is a natural phenomenon that affects domestic animals in tropical, sub-tropical and often in temperate regions of the world during summer months. Heat and humidity during the summer months combine to make an uncomfortable environment for dairy cattle. Heat stress negatively impacts a variety of dairy parameters resulting in economic losses ^33^. Response to heat stress varies with species and genetic groups within species ^5,34,35^. In this study, transcriptome of genetic groups – Vrindavani and Tharparkar cattle under heat stress was evaluated to understand their differential response to heat stress.

Animals (n=5) of both the genetic groups were exposed to a temperature of 42 °C for 7 days. Around 5^th^−6^th^ day, short term heat acclimation occurs ^30,31^. This time point was selected to understand the differences in systems biology to heat stress in the two genetic groups. Initially, heat stress indicators - RR, RT, and T3 level were evaluated. RR was found to increase in both genetic groups under heat treatment and the increase in Vrindavani was found to be significantly (P<0.05) different from that in Tharparkar. A positive correlation exists between RR and heat treatment ^36-38^. This increase is an attempt to dissipate excess body heat by vaporizing more moisture in expired air or response to a greater requirement of oxygen by tissues under heat stress. Also, the physiological response to heat stress includes reduced heat production, which is achieved by lowering feed intake and thyroid hormone secretion ^39^. T3 level increases under heat stress ^40,41^. A significant increase in T3 level in Vrindavani as compared to Tharparkar indicates an effective regulatory mechanism in modulating T3 levels in Tharparkar in response to heat stress. The T3 triggered metabolism may be one of the reasons that increases heat production resulting in high rectal temperature in Vrindavani in comparison to Tharparkar as was found in our study. The significant increase in RR, RT and T3 level in Crossbreed than in Tharparkar, suggests the inability of Vrindavani to cope up with heat stress in comparison to Tharparkar.

A phenotype is defined by the changes in systems biology. Transcriptome profiling by RNA-seq is the most common methodology to study the changes in systems biology. RNA profiling based on next-generation sequencing enables to measure and compare gene expression patterns^21^. The transcriptome of Tharparkar and Vrindavani indicated differential response to heat stress as evident from the DEGs, which were either distinct to both or have a difference in expression. The number of DEGs were higher in Vrindavani than in Tharparkar, suggesting a greater dysregulation in systems biology in Vrindavani. Among the dysregulated genes, the number of upregulated genes were more than the downregulated genes in both genetic groups. However, a contrast in expression was observed with 18.5 % of upregulated genes in Vrindavani, were downregulated in Tharparkar and 17.5% upregulated genes in Tharparkar were downregulated in Vrindavani.

The differentially expressed genes in each genetic group were functionally annotated using both reactome and IPA. In the reactome, the enriched pathways were identified and in IPA, based on the expression of the DEGs the activated/inactivated/not-activated pathways were identified. In reactome, on functionally annotating the DEGs of each of the genetic groups, key pathways related to stress – neutrophil degranulation ^42^, L13a-mediated translational silencing of ceruloplasmin expression ^43^, ubiquitination and proteasome degradation ^44^, SRP-dependent cotranslational protein targeting to membrane ^45^, Nonsense-Mediated Decay ^46^ and selenocysteine synthesis ^47^ were enriched in both the genetic groups. Among the common genes, the genes upregulated in Tharparkar but downregulated in Vrindavani showed enrichment towards cellular response to stress besides the above-mentioned pathways. This indicated that Tharparkar must be responding to withstand heat stress. Further, on evaluating the common genes that were upregulated in Vrindavani and downregulated in Tharparkar, pathways - chromatin modifying enzymes, chromatin organization and death receptor signaling indicated towards apoptosis and DNA repair in Vrindavani. These findings were further corroborated with the pathway enrichment analysis of unique genes in both the genetic groups. IPA analysis of disease and biological function showed repair of DNA and ubiquitination, activated in Vrindavani and senescence and degranulation of cells inactivated in Tharparkar. The results from reactome and IPA suggested Tharparkar may be more resilient than Vrindavani.

IPA revealed activation or inactivation of several pathways in both the genetic groups. It is known that - EIF2 signalling, helps in initiation of global protein translation ^48^; MAPK-signalling pathway, induces cell proliferation ^49^; androgen signalling, enhances pro-survival and anti-apoptotic activity in cell ^50^; a-Adrenergic signalling, maintains immune defence mechanism ^51^ and, helps in tissue repair upon stress ^52^ and increases angiogenesis ^53^; integrin pathway, resists the cell against apoptosis and other environmental insults ^54^; IL-3 signalling, aids in cell survival and haematopoiesis ^55^; CXCR4 signalling modulates cell survival and cell motility ^56^ and ; Phospholipase C signalling aids in cell survival in stress through protein kinase C dependent phosphorylation of BCL-2 ^57^. Inactivation of these pathways except MAPK-signalling pathway in Vrindavani and activation of α-Adrenergic signalling, Androgen signalling, EIF2 signalling and MAPK signalling in Tharparkar indicates that the systems biology in Tharparkar is moving towards countering the effects due to heat stress.

On correlating the reactome data with the IPA four major physiological processes - elicitation of unfolded protein response (UPR) in cells; Ubiquitination; Induction of apoptosis and; Imbalance in production of ROS and antioxidants were considered for further evaluation (Figure 8). Heat shock and its associated genes are involved in elicitation of unfolded protein response (UPR) in cells. Most of the heat shock genes were found upregulated in Tharparkar and downregulated in Vrindavani. The increased HSP levels have been found positively correlated with tolerance in many species^58,59^. HSF1, that positively regulates the transcription of *HSP70* and *HSP90* ^60,61^ was found upregulated in Tharparkar and downregulated in Vrindavani. Upregulation of *HSF1, HSP70 and HSP90* in Tharparkar and vice-versa in Vrindavani corroborates to state that Tharparkar is better equipped to counter heat stress than Vrindavani. Further, to ensure that the HSP70 in Tharparkar is maintained at an optimum level, dysregulation of *CAMK2D* and *GSK3A* seems to act as negative feedback. CAMK2D that induces the transcription of HSP70 via HSF1 ^62^ was found downregulated in Tharparkar. GSK3A that inhibits the trimerization of HSF1 that is needed for the induction of HSP70 ^63^ was found upregulated in Tharparkar. The decreased level of HSP70 in Vrindavani makes it inevitable that such negative feedbacks would further reduce its level and hence, *GSK3A* was found downregulated and *CAMK*, upregulated (Figure 8).

**Figure 8:**
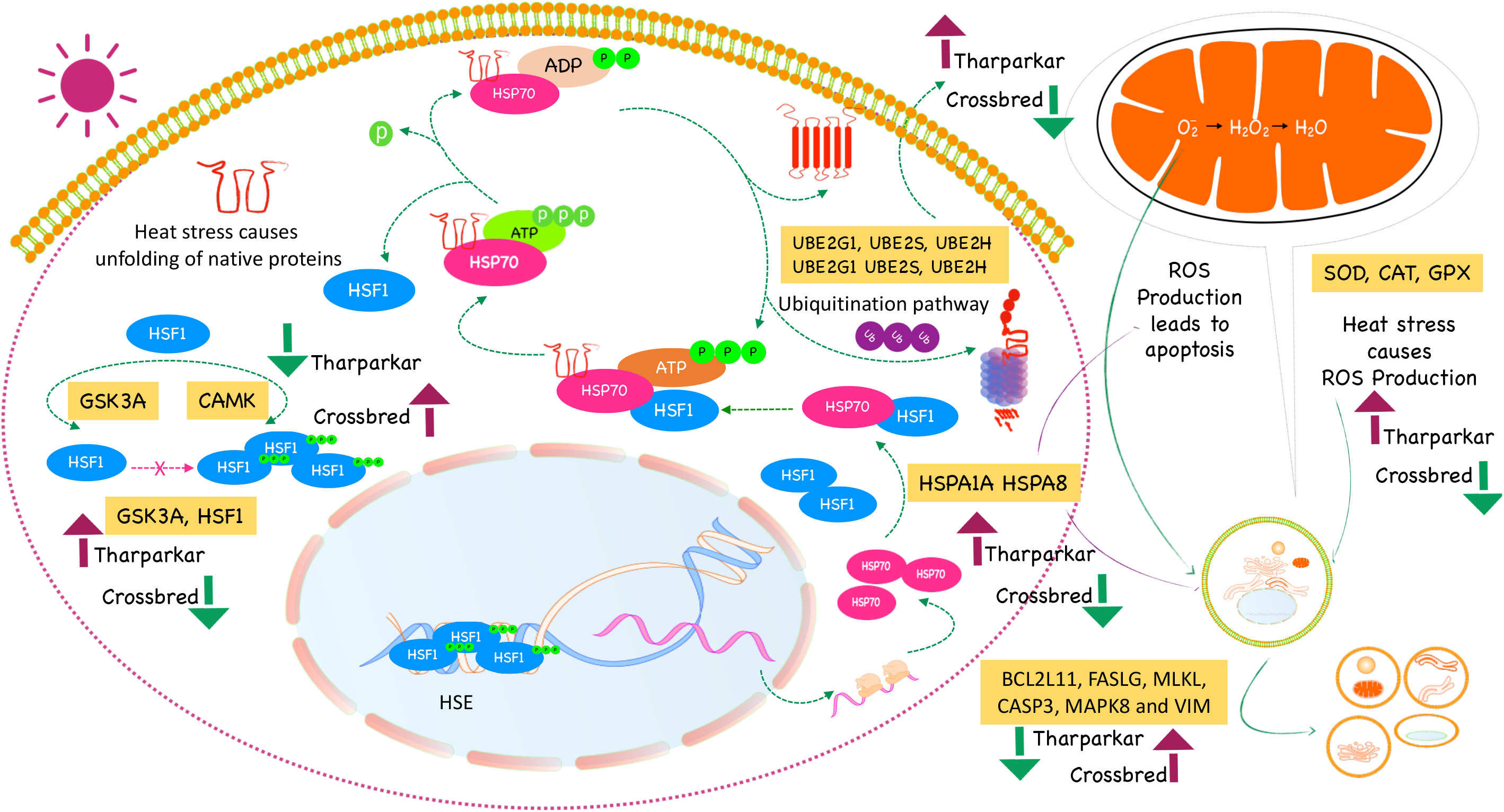
Predicted interplay of molecules that is underway during heat stress in both groups: Heat stress causes unfolding of native proteins. HSP70 acts as a chaperone to facilitate refolding to restore the structure of unfolded proteins. Under normal condition, HSP70 is bound to HSF1 thereby preventing HSF1 to promote transcription of HSP70. Under heat stress ATP binds to the HSP70 and HSF1 complex to release HSF1, promoting the binding of the unfolded protein to HSP70 and ATP. CAMK2D that induces the transcription of HSP70 via HSF1 was found downregulated in Tharparkar. GSK3A that inhibits the trimerization of HSF1 that is needed for the induction of HSP70 expression was found upregulated in Tharparkar. The decreased level of HSP70 in Vrindavani makes it inevitable that such negative feedbacks would further reduce its level and GSK3A was found downregulated and CAMK2D, upregulated. Further, in Tharparkar, HSP70 tends to activate ubiquitination pathway to decrease the accumulation of unfolded proteins that can’t be refolded by it. This pathway activation is supported by upregulation of E3 ligases (UBE2G1, UBE2S, and UBE2H) in Tharparkar. However, the E3 ligase in Vrindavani was found downregulated. With HSP70 being upregulated and having cytoprotection activity, Tharparkar shows the decline in apoptosis as compared to Vrindavani. This is supported by downregulation of BCL2L11, FASLG, MLKL, CASP3, MAPK8 and VIM in Tharparkar and vice-versa. Besides, higher expression of the antioxidants (SOD, CAT, GPX) in Tharparkar enables it to cope up with higher levels of free radicals generated as a result of heat stress while Vrindavani is unable to do so. Green arrow indicates downregulation and Maroon arrow indicates upregulation.

Ubiquitination is positively correlated with heat tolerance ^64,65^. Ubiquitin-Proteasome System (UPS) regulates the levels of proteins and acts by removing the misfolded or damaged proteins that may accumulate as a result of exposure to abiotic stress. Malfunctioning of ubiquitin-proteasome system (UPS) could have negative consequences for protein regulation, including loss of function ^66^. In Tharparkar after heat acclimation, HSP70 tends to activate the ubiquitination pathway to minimize the accumulation of the unfolded proteins that can’t be refolded by it ^67^. This pathway activation is supported by upregulation of E2 ligases - *UBE2G1, UBE2S*, and *UBE2H* that catalyze covalent attachment of E2 to E3 ^68-71^ in Tharparkar. USP7 that deubiquitinates target proteins ^72,73^ was found upregulated in Vrindavani and downregulated in Tharparkar. Further, a group of molecules – *VCP, SERP1*, and *CALR* that ensure the protection of naïve proteins during their transport within the cell ^74-76^ were found upregulated in Tharparkar and downregulated in Vrindavani. Unlike Vrindavani, Tharparkar is not only endowed with higher expression of the scavengers of misfolded proteins but also with protectors of naïve unfolded proteins.

Activation of apoptosis pathway is one of the major physiological processes linked with heat stress. Among the apoptotic genes, *BCL2L11, FASLG, MLKL, CASP3, MAPK8*, and *VIM* have been found upregulated in Vrindavani and downregulated in Tharparkar under heat stress. BCL2L11 induces apoptosis by neutralizing key molecules of pro-survival BCL2 sub-family ^77,78^, FASLG transduces the apoptotic signal into cells^79,80^, CASP3 activates caspases and executes apoptosis ^81^, and MAPK8, MLKL, and VIM also induce apoptosis ^82,83^. PTEN signaling pathway that drives apoptosis ^84,85^ was found inactivated in Tharparkar and activated in Vrindavani. This indicates a relatively higher probability of apoptosis in Vrindavani than in Tharparkar.

The ability to balance the ROS and antioxidant level, is one of the key factors that would determine the tolerance of an individual to heat stress. The antioxidant triad of GPX, SOD, and CAT that forms the first line of defense against reactive oxygen species ^86-88^, was found upregulated in Tharparkar and downregulated in Vrindavani. Additionally, genes belonging to Peroxiredoxins - *PRDX3, PRDX5* and *PRDX6* that catalyze the reduction of hydrogen peroxide and organic hydroperoxides ^89-93^,were also found upregulated in Tharparkar and were either downregulated or not-differentially expressed in Vrindavani. Higher expression of the antioxidants in Tharparkar enables it to cope up with higher levels of free radicals generated as a result of heat stress while Vrindavani is unable to do so.

## Conclusion

A contrast in expression was observed with 18.5 % of upregulated genes in Vrindavani were downregulated in Tharparkar and 17.5% upregulated genes in Tharparkar were downregulated in Vrindavani. Transcripts of molecules that stimulate heat shock response, Ubiquitination, unfolded protein response and antioxidant level were found upregulated in Tharparkar and downregulated in Vrindavani. EIF2 Signaling that promotes protein translation and PTEN signaling that drives apoptosis were found activated and inactivated in Tharparkar, respectively and vice-versa in Vrindavani. We found relevant molecules/genes dysregulated in Tharparkar in the direction that counters heat stress. A proposed contrasting interplay of molecules in both the two groups is shown in Figure 8. To the best of our knowledge this is a comprehensive comparison between Tharparkar and Vrindavani at a global level using transcriptome analysis.

## Methods

### Experimental condition and Ethical Statement

The animals used for the study were from the Indian Veterinary Research Institute. The permission to conduct the study was granted by Indian Veterinary Research Institutional Animal Ethics Committee (IVRI-IAEC) under the Committee for Control and Supervision of Experiments on Animals (CPCSEA), India, vide letter no 387/CPSCEA. Genetic groups - Tharparkar (Indigenous breeds) and Vrindavani (synthetic Crossbred) were considered in this study. Prior to experiment, the animals – 05 Tharparkar and 05 Vrindavani cattle, were acclimatized for 15 days outside the Psychometric chamber. The experiment was conducted during October when the environmental Temperature Humidity Index (THI) was 73.0242. These animals were exposed in Psychometric chamber at 42 °C for six hours for 7 days (THI =78.5489). All the animals were fed with wheat straw and concentrate mixture in 60:40 ratios. Respiration rate (RR) and rectal temperature (RT) of animals from each genetic group were measured on 0 day (Control, n=5) before exposure to Psychometric chamber and on 7^th^ day of heat exposure (Treated, n=5). Blood samples were collected from the animals at the mentioned time points and serum concentration of Triiodothyronine (T3) was estimated by RIA technique using T_3_ ^125^I (Immunotech) as per the manufacturer’s instructions.

### RNA sequencing (RNA-seq)

PBMCs were collected from the blood samples using Ficol histopaque gradient method. Total RNA from each of the collected samples (PBMCs) was isolated using the RNeasy Mini kit (Qiagen GmbH, Germany) according to the manufacturer’s protocol. The integrity and quantity of isolated RNA were assessed on a Bioanalyzer 2100 (Agilent Technologies, Inc). The library was prepared using NEBNext Ultra RNA Library Prep Kit for Illumina (NewEngland Biolabs Inc.) following the manufacturer’s protocol. Approximately, 100ng of RNA from each sample was used for RNA library preparation. The quality of the libraries was assessed on Bioanalyzer. Libraries were quantified using a Qubit 2.0 Fluorometer (Life technologies) and by qPCR. Library (1.3ml, 1.8pM) was denatured, diluted and loaded onto a flow cell for sequencing. cDNA library preparation and Illumina Sequencing was performed at Sandor Life Sciences Pvt. (Hyderabad, India). Finally, the RNA-seq data were provided in FASTQ format.

### Data processing

The sequenced reads were paired-end and 150bp in length. Quality control checks on raw sequence data from each sample were performed using FastQC (Babraham Bioinformatics). Processing of the data was performed using prinseq-lite software ^94^ to remove reads of low quality (mean phred score 25) and short length (< 50) for downstream analysis. The data was submitted to the GEO database with accession number GSE136652.

### Identification of Differentially Expressed Genes (DEGs)

*Bos taurus* reference genome (release 94) and its associated gene transfer file (GTF) were downloaded from Ensembl FTP genome browser ^95^. The reference genome was prepared and indexed by RNA-Seq by expectation maximization (RSEM) ^96^ by rsem-prepare-reference command. Further, the clean reads obtained from filtering of raw data were aligned to the indexed reference genome by Bowtie2 ^97^ to estimate gene abundance in counts by rsem-calculate-expression command. To compare the gene expression levels among different samples, the aligned reads were used to generate a data matrix by rsem-generate-data-matrix command. In each genetic group, all the samples of day 7 (treated) were compared with the day 0 (Control) for the calculation of differential gene expression by edgeR ^98^ package. The Ensemble IDs of the differentially expressed genes (DEGs) were converted to the respective gene ID by g: Convert of g: Profiler ^99,100^.

### Functional Analysis of DEGs - Reactome and Ingenuity pathway analysis (IPA)

The differentially expressed genes in both the genetic groups were functionally annotated in Reactome ^101^ and IPA (Ingenuity Pathway Analysis). Reactome provides bioinformatics tools for visualisation, interpretation and analysis of pathway knowledge to support basic research. Reactome analysis was done for all DEGs in both genetic groups, common DEGs and for unique DEGs for each genetic group. For common DEGs, analysis was also done for contrasting genes (genes upregulated in Tharparkar but downregulated in Vrindavani and the vice-versa).

QIAGEN’s IPA (QIAGEN, Redwood City, USA) ^102^ is used to quickly visualize and understand complex omics data and perform insightful data analysis and interpretation by placing experimental results within the context of biological systems. Here, IPA was used to analyze the identified DEGs of Vrindavani and Tharparkar. The list of DEGs from each genetic group was used to identify the canonical pathways and the most significant biological processes against Ingenuity Pathways Knowledge Base (IKB). Core analysis for each dataset was performed to know activated (Z score > 2) or inactivated (Z score < −2) canonical pathways. Upstream regulators - Transcription factors and microRNAs were identified. Significant Diseases and functions were also identified.

The results obtained from Reactome and IPA were correlated to narrow down to processes - Induction of apoptosis, Ubiquitination, elicitation of unfolded protein response (UPR) in cells and Imbalance in production of ROS and antioxidants. The unique genes involved in all related pathways to the processes mentioned above were extracted from the Reactome analysis and were found to be 246 in number. From among these genes the number of differentially expressed genes in Tharparkar and Vrindavani, and the common genes between the genetic groups were extracted.

### Predicted protein-protein interaction network

Protein interaction network (interactome) analysis provides an effective way to understand the interrelationships between genes ^103^. Protein-protein interactions (PPI) among the common genes involved in the processes mentioned, were retrieved using String database ^104^. The degree was calculated using igraph package (https://cran.r-project.org/web/packages/igraph/index.html). The PPI network was then visualized using Cytoscape software V. 3.7 ^105^ using DEGs and degree value from igraph.

### Validation of reference genes identified

The quantity of data generated from RNA sequencing is large and therefore it is important to validate differential expression by real-time RT-PCR. Genes - *BCL2L11, FASLG, CASP3, CAT, SOD1, GSK3A, CALR, HSF1, APC*, and *GPX3* were selected based on their role in heat stress and qRT-PCR was performed on Applied Biosystems 7500 Fast system. *GAPDH* was taken as the internal control. Each of the samples was run in triplicates and relative expression of each gene was calculated using the 2^−ΔΔ*CT*^ method with control as the calibrator ^106^.

### Statistical Analysis

Respiration rate, Rectal temperature and T3 level were compared using student’s *t*-test in JMP9 (SAS Institute Inc., Cary, USA) to test the significance of the difference between the control (0 day) and treated (7^th^ day). This comparison was done within and between genetic groups. Differences within/between groups were considered significant at *P* ≤ 0.05.

## Declarations

### Ethics approval and consent to participate

The permission to conduct the study was granted by Indian Veterinary Research Institutional Animal Ethics Committee (IVRI-IAEC) under the Committee for Control and Supervision of Experiments on Animals (CPCSEA), India, vide letter no 387/CPSCEA.

### Consent for publication

Not applicable.

### Availability of data and materials

The data was submitted to the GEO database with accession number GSE136652

### Authors’ contributions

AKT and RG conceived and designed the research. SG, SmS, AS,AV, VV, PK, ShS and GS conducted the wet lab work. RINK, ARS, NH, WAM, MRP, SK, AP and RG analyzed the data. RINK, ARS, MRP, RG, AS and GS helped in manuscript drafting and editing. AKT and RG proofread the manuscript

## Acknowledgements

We also thank Department of Biotechnology, Govt of India for providing fellowship and contingency for RK (DBT Fellow No. DBT/2017/IVRI/768), ARS (DBT Fellow No. DBT/2014/IVRI/170) and WAM (DBT Fellow No. DBT/2017/IVRI/769).

## Legends

**Supplementary.Figure 1:** Comparison of activated/inactivated pathways in Vrindavani and Tharparkar. Activated pathways have Z score > 2 and indicated by red colour while inactivated pathways are having Z score < − 2 and indicated by green colour.

**Supplementary Figure 2:** Comparison of activated/inactivated miRNAs in Vrindavani and Tharparkar as predicted by IPA upstream analysis. Activated pathways have Z score > 2 and indicated by red colour while inactivated pathways are having Z score < − 2 and indicated by green colour.

**Supplementary Figure 3:** Comparison of activated/inactivated Transcription factors as predicted by IPA upstream analysis (Transcription factors of Vrindavani are red-coloured and Tharparkar are blue-coloured) vis-à-vis their Log2FC in both genetic groups. The encircled ones are common to both groups.

**Supplementary Figure 4:** Predicted Protein-protein interaction network of expressed genes common to Tharparkar and Vrindavani. The diameter of the node represents the connectivity/degree of the node among the genes.

**Supplementary Figure 5:** Validation of RNA sequencing data by Real-Time data in Vrindavani (a) and Tharparkar (b). The expression of 10 selected genes was found in concordance with RNA Sequencing data. The correlation (r2 = 0.9942 in (a) and 0.9972 in (b)) was found to be significant (P< .01) in both cases.

