## Supplementary figures and images for "Systems Biology under heat stress in Indian Cattle"

### Supplementary Figure 1

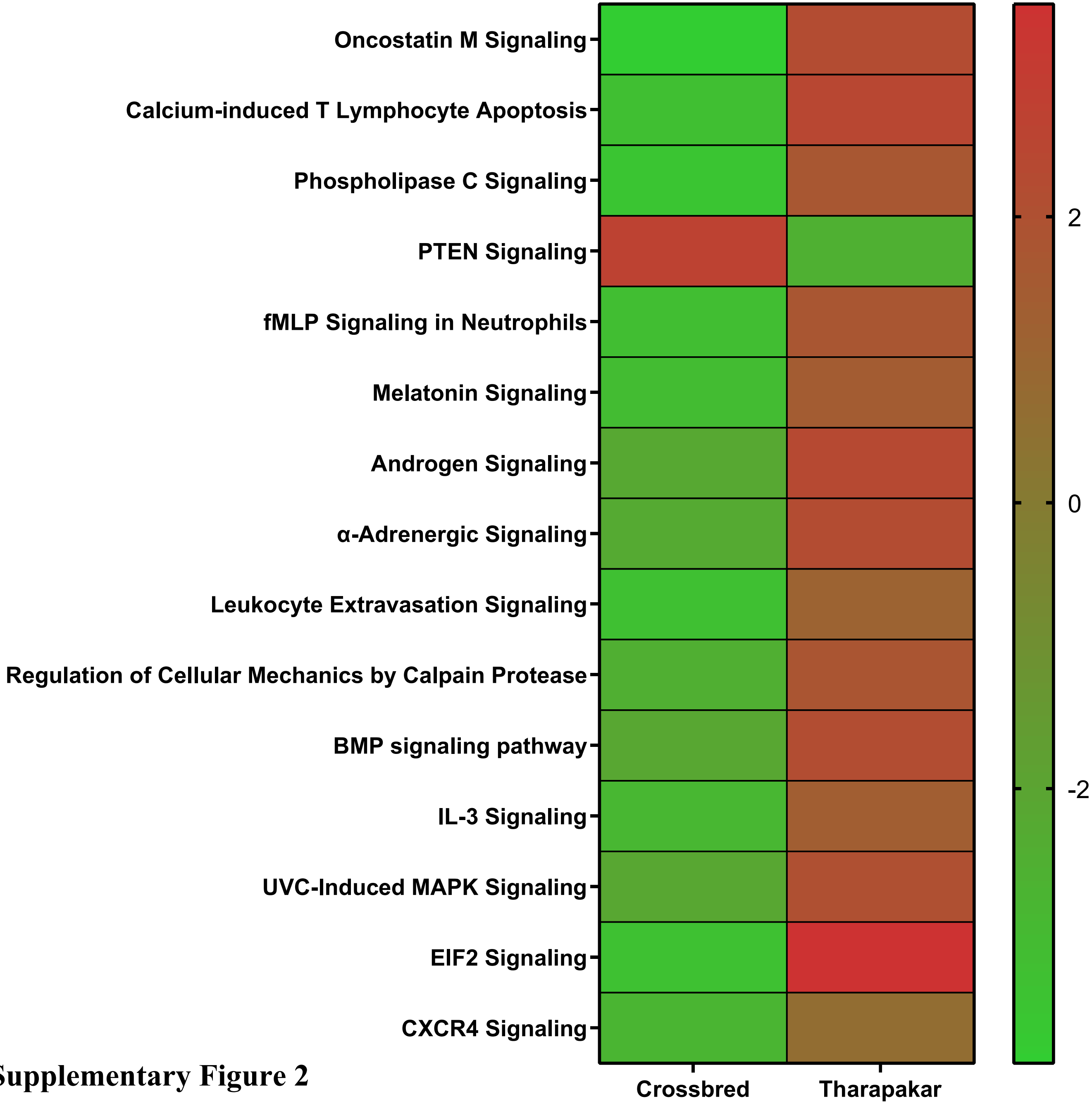


**Supplementary Figure 2**

### Supplementary Figure 2

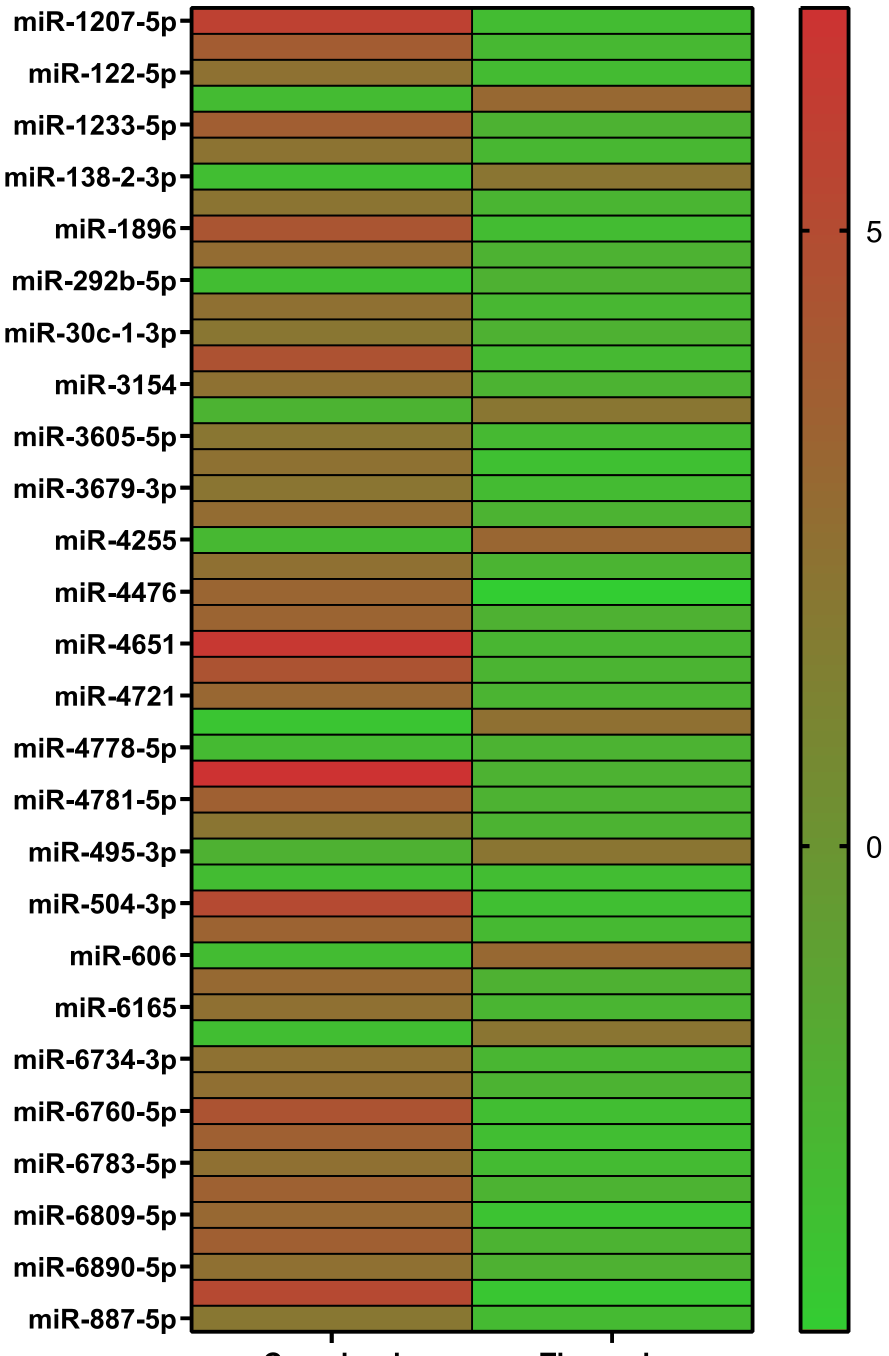


Supp Figure 3

### Supplementary Figure 3

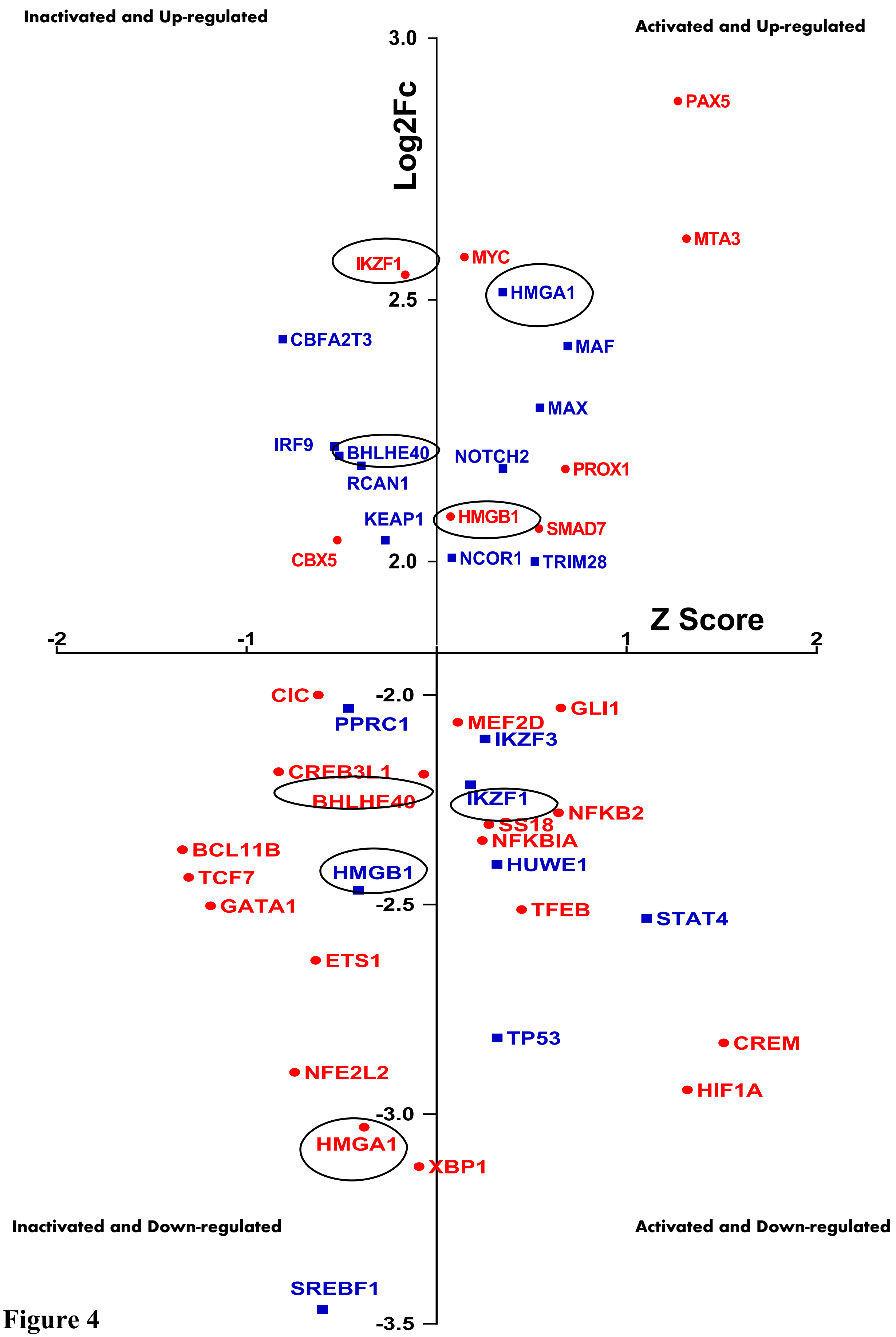


Supplemenatary Figure 4

### Supplementary Figure 4

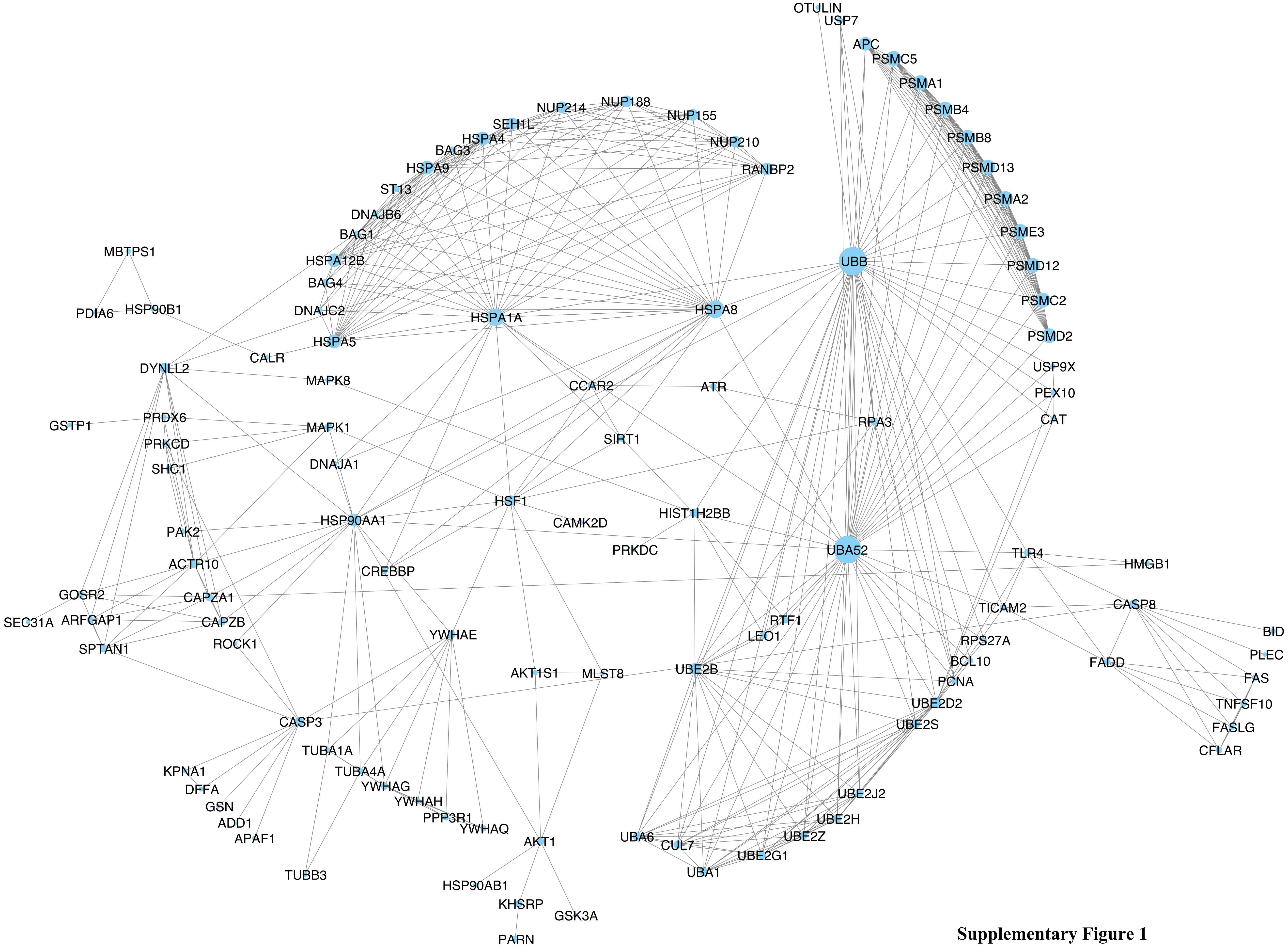


**Supplementary Figure 1**

### Supplementary Figure 5

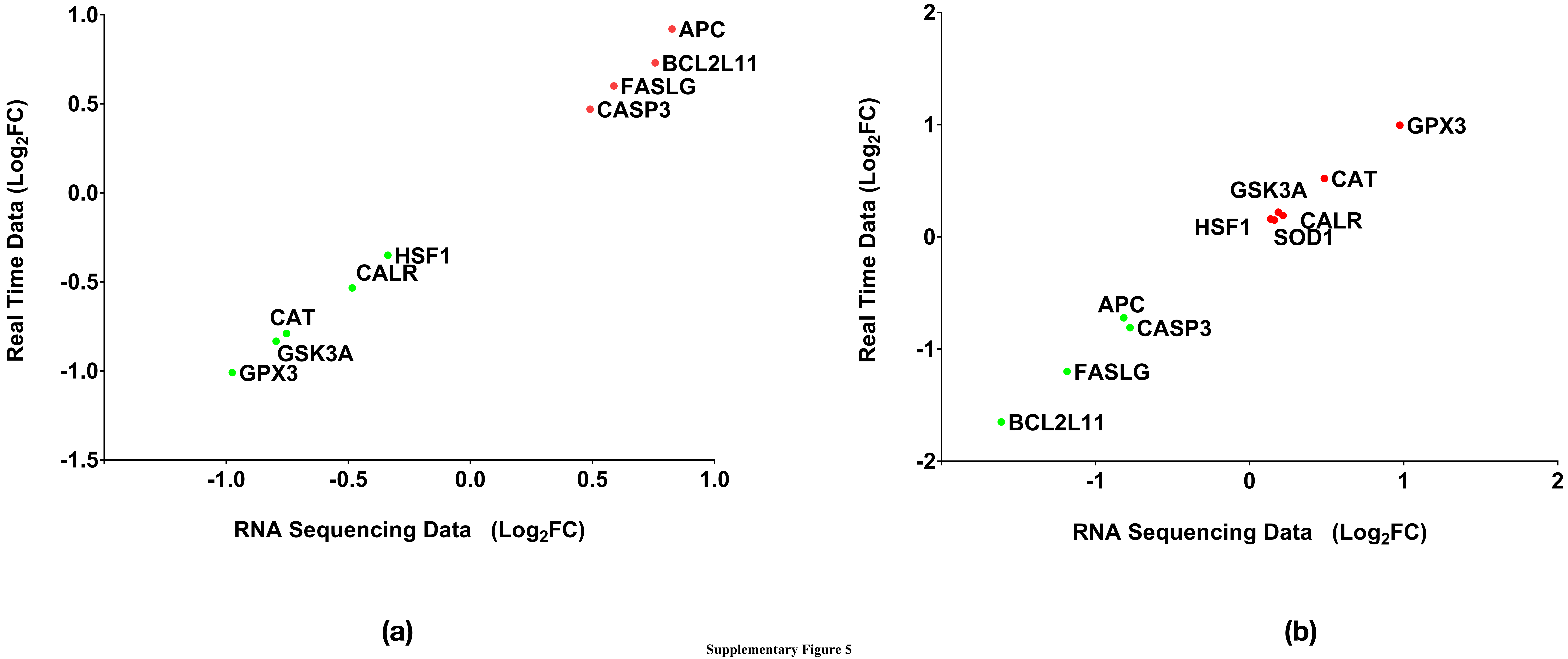


Supplementary Figure 5
