## Supplementary Table 2 for "Systems Biology under heat stress in Indian Cattle"

Supplementary Table 2: $log_2FC$ from RNAseq and FC from qRT-PCR of two genetic groups- Vrindavani and Tharparkar

| Gene | $log_2FC$ from RNAseq (Vrindavani) | FC from qRT-PCR (Vrindavani) | $log_2FC$ from RNAseq (Tharparkar) | FC from qRT-PCR (Tharparkar) |
|---|---|---|---|---|
| BCL2L11 | 0.757638 | 0.65 | -1.61229 | -1.65 |
| FASLG | 0.588103 | 0.6 | -1.18454 | -1.2 |
| CASP3 | 0.490952 | 0.53 | -0.77592 | 0.81 |
| CAT | -0.753080282 | -0.79 | 0.485014 | 0.52 |
| SOD1 | | | 0.137292 | 0.16 |
| GSK3A | -0.795474096 | -0.833 | 0.186747 | 0.22 |
| CALR | -0.483691773 | -0.534 | 0.161541 | 0.15 |
| HSF1 | -0.337490189 | -0.35 | 0.217194 | 0.19 |
| APC | 0.826575 | 0.92 | -0.81711 | -0.721 |
| GPX3 | -0.975291898 | -1.01 | 0.975640381 | 0.94 |
